# 12-Lipoxygenase inhibition improves glucose homeostasis and obesity-associated inflammation in human gene replacement mice

**DOI:** 10.1101/2025.01.10.632274

**Authors:** Kerim B. Kaylan, Titli Nargis, Kayla Figatner, Jiayi E. Wang, Sarida Pratuangtham, Advaita Chakraborty, Isabel Casimiro, Jerry L. Nadler, Matthew B. Boxer, David J. Maloney, Ryan M. Anderson, Raghavendra G. Mirmira, Sarah A. Tersey

## Abstract

Obesity-associated inflammation is characterized by macrophage infiltration into peripheral tissues, contributing to the progression of prediabetes and type 2 diabetes (T2D). The enzyme 12-lipoxygenase (12-LOX) catalyzes the formation of pro-inflammatory eicosanoids and is known to promote the migration of macrophages, yet its role in obesity-associated inflammation remains incompletely understood. Furthermore, differences between mouse and human orthologs of 12-LOX have limited efforts to study existing pharmacologic inhibitors of 12-LOX. In this study, we utilized a human gene replacement mouse model in which the gene encoding mouse 12-LOX (*Alox15*) is replaced by the human *ALOX12* gene. As a model of obesity and dysglycemia, we administered these mice a high-fat diet. We subsequently investigated the effects of VLX-1005, a potent and selective small molecule inhibitor of human 12-LOX. Oral administration of VLX-1005 resulted in improved glucose homeostasis, decreased β cell dedifferentiation, and reduced macrophage infiltration in islets and adipose tissue. Analysis of the stromal vascular fraction from adipose tissue showed a reduction in myeloid cells and cytokine expression with VLX-1005 treatment, indicating decreased adipose tissue inflammation. In a distinct mouse model in which *Alox15* was selectively deleted in myeloid cells, we observed decreased β cell dedifferentiation and reduced macrophage infiltration in both islets and adipose tissue, suggesting that the effects of VLX-1005 may relate to the inhibition of 12-LOX in macrophages. These findings highlight 12-LOX as a key factor in obesity-associated inflammation and suggest that 12-LOX inhibition could serve as a therapeutic strategy to improve glucose homeostasis and peripheral inflammation in the setting of obesity and T2D.

## INTRODUCTION

Obesity results from prolonged nutrition excess and diets rich in saturated fats. Even without impaired glucose tolerance or type 2 diabetes (T2D), individuals can develop hyperinsulinism and other features of dysregulated insulin action, such as peripheral insulin resistance (1,2). Hyperinsulinism is the adaptive result of increased peripheral demand for insulin secretion from islet β cells in the setting of insulin resistance (3,4). However, in some cases, the ability of β cells to adaptively compensate is limited by many factors, including oxidative stress, glucolipotoxicity, endoplasmic reticulum (ER) stress, and infiltration of the islet by inflammatory macrophages (5,6). Additionally, infiltration of adipose tissue by macrophages and other immune cells leads to inflammation and the systemic release of proinflammatory adipokines, worsening insulin resistance and systemic inflammation (7,8).

Signaling via the products of 12-lipoxygenase (12-LOX) enzymes has previously been implicated in the progressive changes associated with obesity and T2D described above (9). As a family, the LOX enzymes facilitate the oxygenation of polyunsaturated fatty acids to form eicosanoids, which are inflammatory lipid intermediates known to contribute to oxidative stress. The eicosanoids produced by 12-LOX enzymes include the proinflammatory metabolite 12-hydroxyeicosatetraenoic acid (12-HETE). The main 12-LOX enzyme that produces 12-HETE in β cells and macrophages in mice is encoded by the *Alox15* gene (9). Whole-body knockout (KO) of *Alox15* in mice protects against worsening of glucose and insulin tolerance after administration of a high-fat diet (HFD), while also improving systemic inflammatory markers and adiponectin secretion (10,11). Pancreas-specific KO of *Alox15* results in complete protection from worsened glucose and insulin tolerance from HFD, greater adaptive islet hyperplasia, and improved glucose-stimulated insulin secretion (12). Consistent with distinct roles of 12-LOX, pancreas-specific *Alox15* KO mice showed no difference in adipose tissue inflammation, suggesting differential roles for 12-LOX in β cells versus macrophages. Lastly, the adipose tissue conditional KO of *Alox15* improves glucose homeostasis and reduces systemic inflammation and macrophage infiltration in adipose tissue in mice treated with HFD (13).

The translational relevance of these findings is unclear, as the relevant LOX enzyme that produces 12-HETE in humans is 12-LOX, which is encoded by *ALOX12*. Human 12-LOX is structurally distinct from its mouse orthologue, and therefore selective inhibitors of human 12-LOX cannot be tested for efficacy in mice (14). VLX-1005 (also known as ML355) is a potent and selective inhibitor of human 12-LOX (15). We have recently shown that oral treatment with VLX-1005 in non-obese diabetic (NOD) mice in which human *ALOX12* replaced mouse *Alox15* delays the onset of autoimmune diabetes and reduces islet inflammation (14). Here, we use the human gene replacement mice on the *C57BL/6J* background (*B6.hALOX12*) to investigate the impact of pharmacologic inhibition of 12-LOX in obesity and dysglycemia. We administered HFD to *B6.hALOX12* mice and evaluated the effect of 12-LOX inhibition with VLX-1005 in the context of both dysglycemia prevention and treatment. We also used complementary animal models, including zebrafish and a macrophage-specific KO of *Alox15*, to understand the impact of VLX-1005 treatment and cell type-specific effects of 12-LOX. Our studies provide evidence that pharmacologic inhibition of 12-LOX may have therapeutic benefits in the setting of obesity and dysglycemia.

## RESULTS

### VLX-1005 treatment improves insulin tolerance in B6.hALOX12 mice with preexisting obesity and glucose intolerance

We first asked if 12-LOX inhibition improves glucose homeostasis when given to mice with preexisting obesity and glucose intolerance. We used VLX-1005, a potent and selective inhibitor of human 12-LOX (15), in a previously described mouse model in which the endogenous mouse *Alox15* gene is replaced by the human *ALOX12* gene on the *C57BL/6J* background (*B6.hALOX12* mice) (14). Prior pharmacokinetic analyses showed sufficient oral bioavailability of VLX-1005 as a spray dried formulation when administered daily at 30 mg/kg to *C57BL/6J* mice (14). Male *B6.hALOX12* mice at 8 weeks of age were first administered a HFD (60% of total calories from fat) for 8 weeks to induce obesity and impaired glucose tolerance. Subsequently, the mice were allocated to receive either vehicle or VLX-1005 (30 mg/kg) by oral gavage (**Figure 1A**). The mice continued to receive both HFD and either vehicle or VLX-1005 for an additional 6 weeks after allocation (14 weeks after starting HFD). Body weight, fat mass, and lean mass of these animals were indistinguishable between treatment groups (**Figure 1B–C**). Random-fed blood glucose measurements showed decreases in the VLX-1005 group at 13 and 14 weeks, although this did not reach statistical significance (**Figure 1D**). Glucose tolerance as measured by glucose tolerance test (GTT) at 12 weeks after starting HFD was not different between the two groups (**Figure 1E**). However, insulin tolerance test (ITT) at 14 weeks after starting HFD showed improved insulin sensitivity in the VLX-1005 group (**Figure 1F**). These data recapitulate the improved insulin sensitivity phenotype seen with whole-body *Alox15* KO mice in which genetic manipulation preceded the onset of obesity and glucose intolerance (10,11).

**Figure 1:**
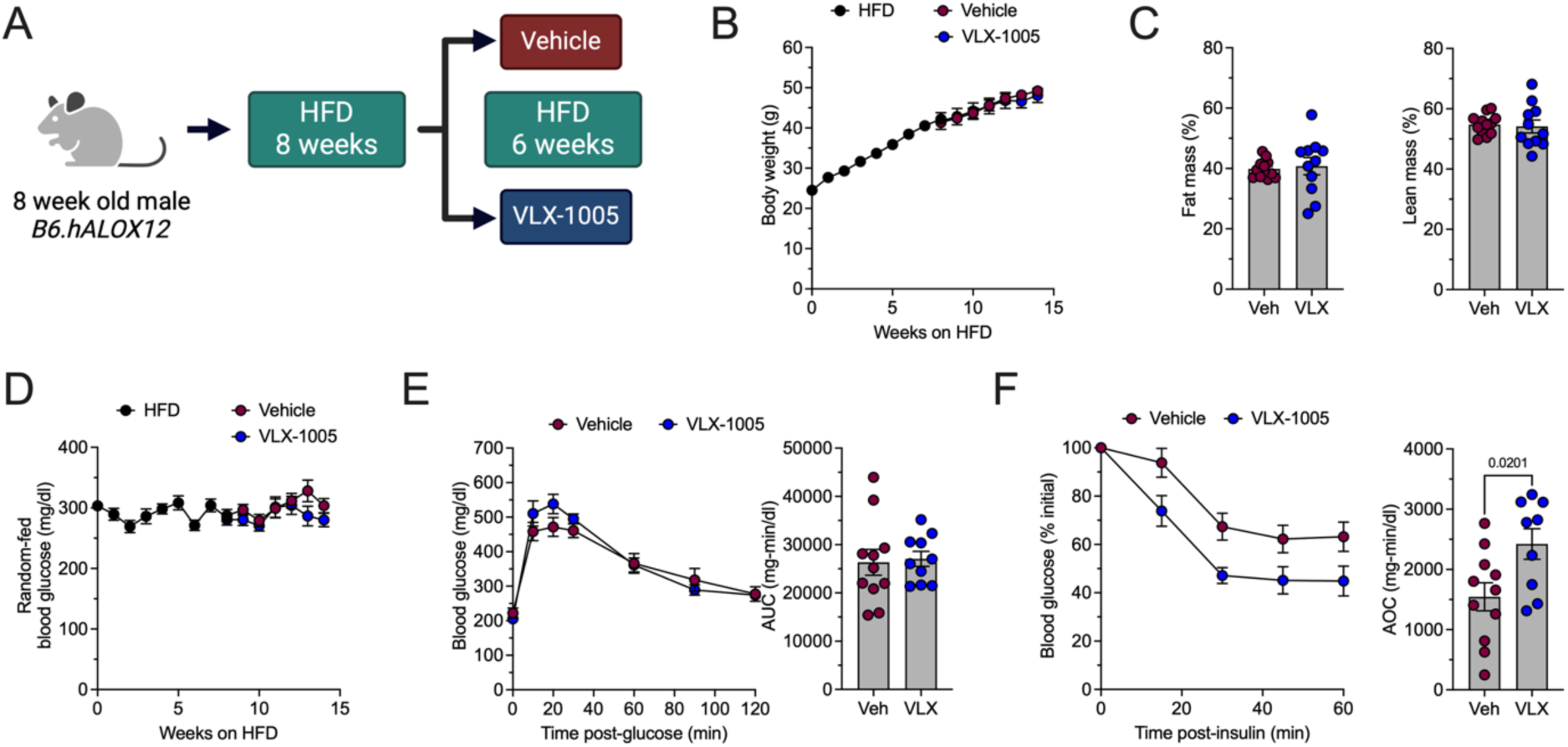
VLX-1005 treatment improves insulin tolerance in B6.h*ALOX12* mice with preexisting obesity and glucose intolerance. (A) Schematic depicting the experimental design. B6.h*ALOX12* mice were treated with HFD for 8 weeks (N=23), followed by addition of either vehicle or VLX-1005 treatment for 6 weeks (N=11-12 per group). (B) Body weight. (C) Fat and lean mass at endpoint. (D) Random-fed blood glucose measurements. (E) Blood glucose measurements from GTT at 12 weeks (*left panel*) with area under the curve (AUC) values (*right panel*). (F) Blood glucose measurements from ITT at 14 weeks (*left panel*) with area over the curve (AOC) values (*right panel*). Data presented as mean with SEM. Single data points represent individual mice. Statistical significance determined by two-tailed Student’s t-testing.

### B6.hALOX12 mice concurrently treated with HFD and VLX-1005 have improved glucose tolerance and insulin sensitivity

We next asked if starting VLX-1005 treatment concurrently with HFD would result in a greater improvement in glucose homeostasis. Male *B6.hALOX12* mice at 8 weeks of age were started on either normal chow diet (NCD, 6% calories from fat) or HFD (60% calories from fat) and concurrently treated with either vehicle or VLX-1005 (30 mg/kg) for 10 weeks (**Figure 2A and Supplemental Figure S1A**). *B6.hALOX12* mice on NCD showed no changes in body weight, fat mass, lean mass, glucose tolerance, or insulin tolerance between the vehicle and VLX-1005 groups (**Supplemental Figure S1B-E**). *B6.hALOX12* mice on HFD exhibited no differences in body weight, fat mass, and lean mass of mice between the vehicle and VLX-1005 groups (**Figure 2B–C**). However, random-fed blood glucose values were lower in the VLX-1005 group as early as 2 weeks after the start of treatment and remained consistently lower throughout the 10 weeks of the experiment (**Figure 2D**). A comparison of the mean random-fed blood glucose in each group showed that the VLX-1005 treatment resulted in a 48 mg/dl lower blood glucose than the vehicle (**Figure 2D**). A GTT at 8 weeks after diet and treatment initiation showed improved glucose tolerance in the VLX-1005 group (**Figure 2E**). Additionally, an ITT at 10 weeks showed slightly increased insulin sensitivity in the VLX-1005 group, although this difference was not statistically significant (**Figure 2F**). Similarly, we observed non-statistically-significant reductions in glucose-stimulated serum insulin levels and homeostatic model assessment for insulin resistance (HOMA-IR), both measures reflective of insulin resistance, in VLX-1005-treated HFD-fed mice compared to vehicle controls (**Figure 2G-H**).

**Figure 2:**
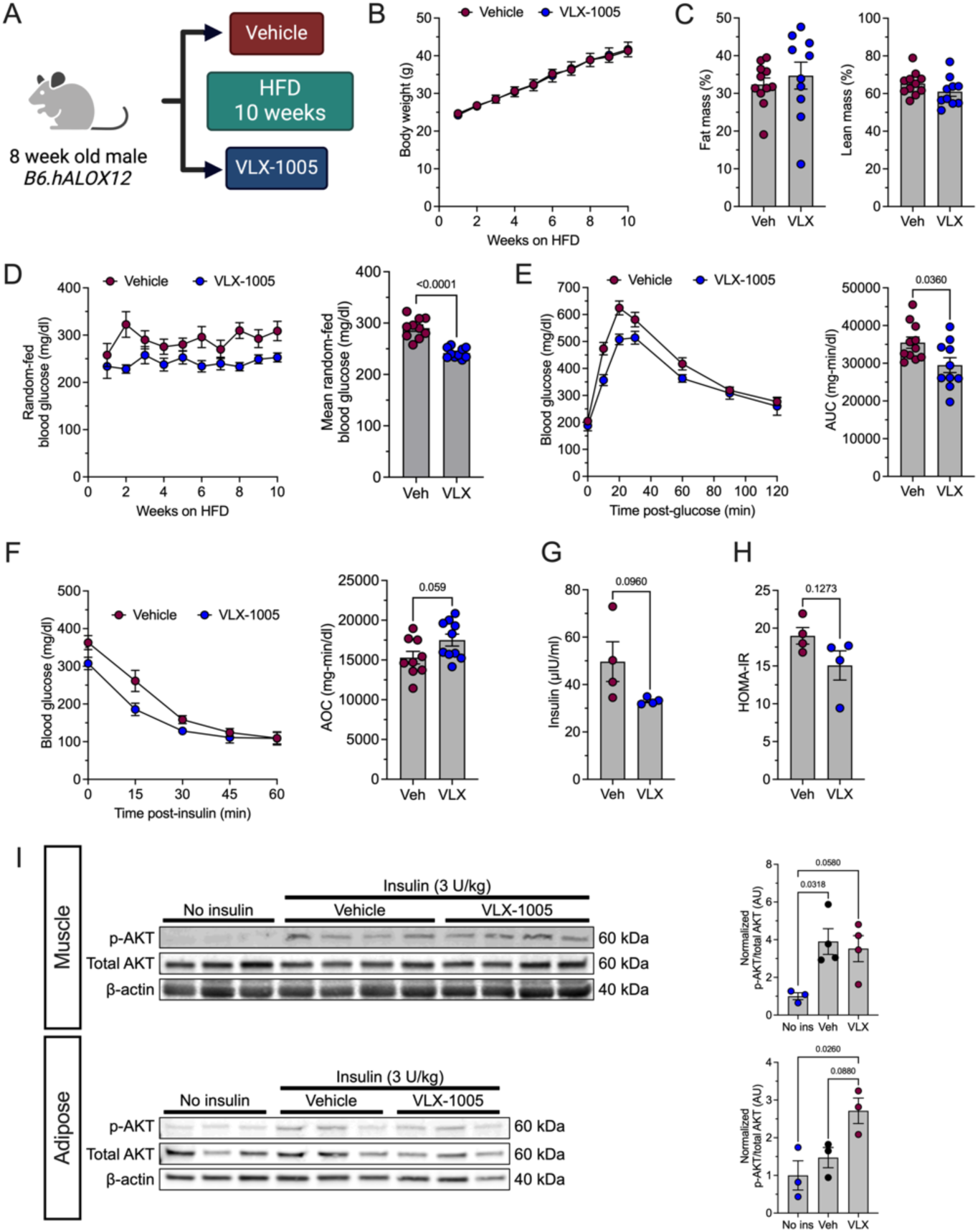
*B6.hALOX12* mice concurrently treated with HFD and VLX-1005 have improved glucose tolerance and insulin sensitivity. (A) Schematic depicting the experimental design. B6.h*ALOX12* mice were treated with HFD concurrently with either vehicle or VLX-1005 for 10 weeks (N=18-19 per group). (B) Body weight. (C) Fat and lean mass at endpoint. (D) Random-fed blood glucose measurements (*left panel*) with per-animal mean over every time point (*right panel*). (E) Blood glucose measurements from GTT at 8 weeks (*left panel*) with AUC values (*right panel*) (N=10-11 per group). (F) Blood glucose measurements from ITT at 10 weeks (*left panel*) with AOC values (*right panel*) (N=9-10 per group). (G) Glucose-stimulated insulin levels at 10 weeks and 10 minutes post-glucose (N=4 per group). (H) HOMA-IR values at 10 weeks (N=4 per group). (I) Western blot analysis in muscle and adipose tissue of phosphorylated AKT (p-AKT) and total AKT (*left panel*) with quantification of the p-AKT/total AKT ratio normalized by β-actin (*right panel*) (N=3-4 per group). Samples of tissue were collected 5 minutes after mice were injected with insulin (3 U/kg). No insulin controls included mice treated with vehicle (right 1 sample) and VLX-1005 (left 2 samples). Data presented as mean with SEM. Single data points represent individual mice. Statistical significance determined by two-tailed Student’s t-testing or one-way ANOVA.

Because physiological measures of insulin sensitivity may not have sufficient sensitivity to capture differences between groups, we next performed a biochemical measurement of tissue insulin sensitivity. Mice were injected with insulin (3 U/kg) 5 minutes prior to sacrifice, and muscle and adipose tissue were collected for analysis of insulin-stimulated phosphorylation of AKT (p-AKT). Immunoblot analysis showed the expected increase in the p-AKT/total AKT ratio with insulin administration but no difference between the vehicle and VLX-1005 treated groups in muscle while adipose tissue showed an increase in p-AKT/total AKT ratio in the VLX-1005 group but not in the vehicle group (**Figure 2I**). These data suggest that inhibition of 12-LOX prior to the onset of obesity improves glucose tolerance and may improve some measures of insulin resistance, especially in adipose tissue.

### VLX-1005 treatment decreases β cell dedifferentiation and macrophage infiltration in islets without changing β cell mass

Next, we examined the effect of 12-LOX inhibition on the pancreatic islets of *B6.hALOX12* mice that had been treated concurrently with (1) NCD and vehicle, (2) HFD and vehicle, or (3) HFD and VLX-1005. While HFD treatment caused the expected increase in β cell mass compared to NCD, mice treated with VLX-1005 did not show differences in β cell mass compared to vehicle controls on HFD (**Figure 3A**). Islet morphology by insulin and glucagon staining appeared grossly similar between the vehicle and VLX-1005 groups on HFD (**Figure 3A**). Because glucose intolerance associated with obesity has been linked to β cell dedifferentiation, we next examined levels of the dedifferentiation marker ALDH1A3 (16,17). By immunostaining quantification, ALDH1A3 was increased with HFD compared to NCD and reduced with VLX-1005 treatment compared to vehicle controls on HFD (**Figure 3B**).

Macrophages infiltrate the islet in the early pathogenesis of obesity and glucose intolerance (18,19). We therefore examined whether treatment with VLX-1005 influences macrophage infiltration into islets. Immunohistochemical staining for the macrophage marker F4/80 showed the expected increased with HFD compared to NCD, and further that VLX-1005 treatment reduced macrophage numbers in islets compared to vehicle controls on HFD (**Figure 3C**). We next asked whether VLX-1005 treatment has a similar effect on macrophage infiltration in a complementary model. We leveraged a transgenic zebrafish line in which macrophages are labeled with green fluorescent protein (GFP) (*Tg(mpeg:eGFP)^gI22^*) (20). Lipid-rich chicken egg has previously been used as a model of metabolic disease in zebrafish larvae, and we adapted a prior protocol used to establish a model of metabolic-associated steatotic liver disease (21,22). We fed *Tg(mpeg:eGFP)^gI22^* zebrafish 5% whole egg solution for 3 hours on days 4- and 6- post fertilization and treated with either vehicle or VLX-1005 (10 µM) starting at 4 days post fertilization (dpf) through endpoint at 7 dpf (**Figure 3D**). Treatment with VLX-1005 reduced macrophages within pancreatic islets to levels similar to controls not treated with 5% whole egg solution (**Figure 3E**). Together, these data show that treatment with VLX-1005 decreases β cell dedifferentiation and macrophage infiltration in islets without associated changes in islet morphology or β cell mass.

**Figure 3:**
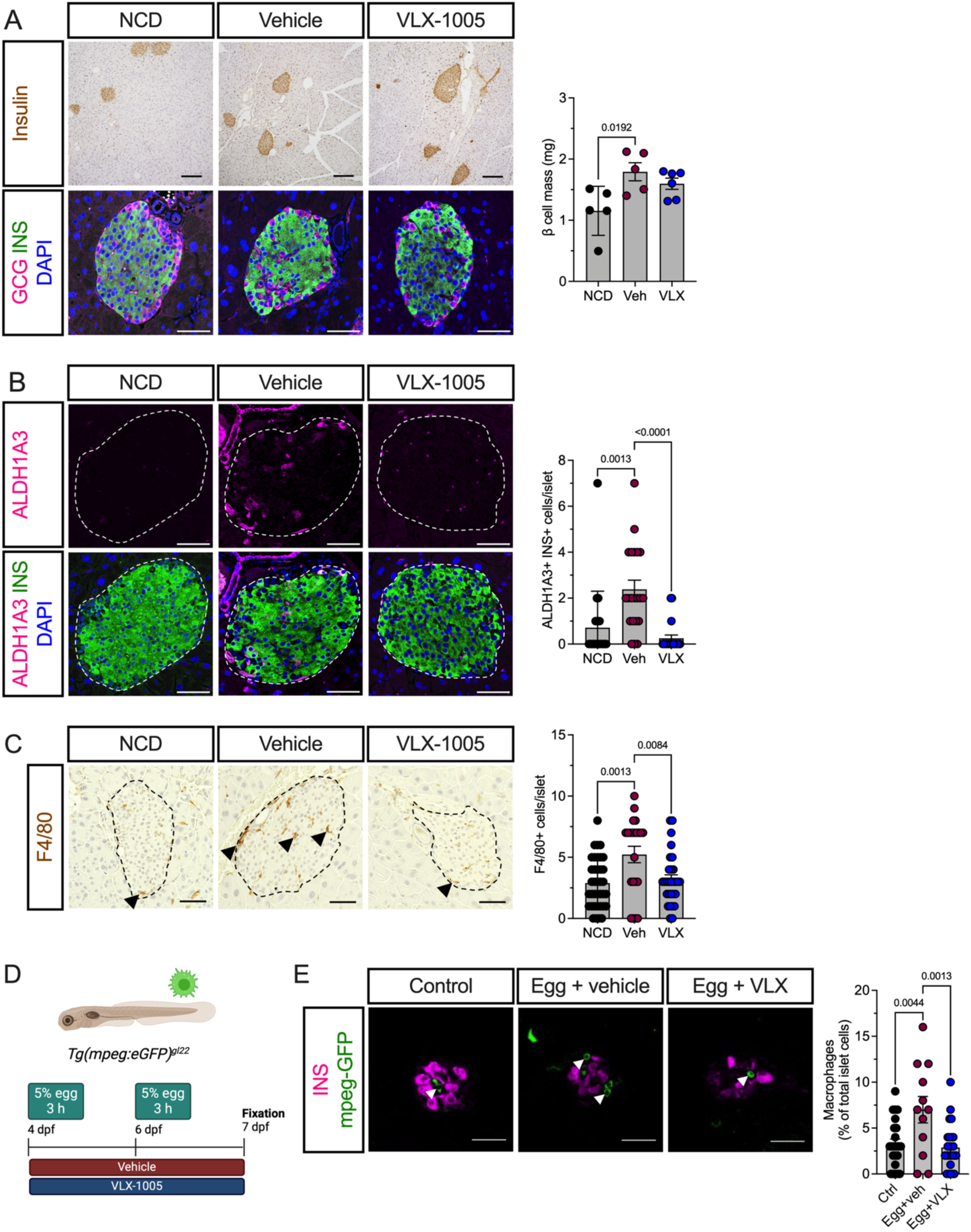
Pancreata of *B6.hALOX12* mice concurrently treated with HFD and VLX-1005 show decreased β cell dedifferentiation and macrophage infiltration in islets without change in β cell mass. (A) Representative images of pancreata from *B6.hALOX12* mice on NCD treated with vehicle or HFD concurrently treated with vehicle or VLX-1005 showing immunohistochemical labeling for insulin (*top left panel*) and immunofluorescence labeling for insulin (green), glucagon (magenta), and nuclei (blue)(*bottom left panel*). Quantification of β cell mass (*right panel*). Upper panel scale bars are 200 µm; lower panel scale bars are 50 µm (N=5-6 per group). (B) Representative images of pancreata with immunofluorescence labeling for ALDH1A3 (magenta), insulin (green), and nuclei (blue)(*left panel*) with quantification of ALDH1A3+ cells per islet (*right panel*). Scale bars are 50 µm. Single data points represent individual islets (N=20-21 islets per group). (C) Representative images of pancreata with immunohistochemical labeling for macrophages with F4/80 (arrowheads) (*left panel*) and quantification of F4/80 cells per islet (*right panel*). Scale bars are 50 µm. Single data points represent individual islets (N=22-38 islets per group). (D) Schematic depicting the design for zebrafish experiments. *Tg(mpeg:eGFP)^gI22^*zebrafish were treated with vehicle or VLX-1005 (10 µM) starting on 4 dpf and 5% whole egg for 3 h on 4 and 6 dpf. Control zebrafish were not treated with 5% egg, vehicle, or VLX-1005. (E) Representative images of zebrafish with labeling for insulin (magenta) and mpeg-GFP (arrowheads, green) (*left panel*). Quantification of macrophages (GFP+) cells as a percentage of total islet cells (*right panel*) (N=12 zebrafish in the control group, N=23-25 zebrafish in the treatment groups). Scale bars are 25 µm. Single data points represent individual islets (N=22-29 islets per group). Data presented as mean with SEM. Statistical significance determined by one-way ANOVA.

### VLX-1005 treatment reduces infiltration of macrophages into adipose tissue in B6.hALOX12 mice

Obesity is also associated with macrophage infiltration in adipose tissue (23,24). Because adipose cell geometry provides information regarding adipose tissue health and increases in adipose cell size are associated with adipose tissue inflammation and insulin resistance (25,26), we next performed immunohistochemistry for F4/80 in the adipose tissue of mice concurrently treated with: (1) NCD and vehicle, (2) HFD and vehicle, or (3) HFD and VLX-1005. We observed increased adipocyte area and qualitative increase in macrophage signal and the number of crown-like structures with HFD treatment (**Figure 4A–B**). Treatment with VLX-1005 resulted in reduced adipocyte area and a qualitative decrease in macrophage signal and the number of crown-like structures in mice compared to vehicle controls on HFD (**Figure 4A–B)**. Using whole-mount immunofluorescence in adipose tissue for macrophage markers F4/80 and iNOS as well as the plasma member marker caveolin to identify adipocytes, we observed reduced counts of proinflammatory macrophages (F4/80+ and iNOS+) with VLX-1005 treatment (**Figure 4C–D**). We next isolated the stromal vascular fraction (SVF) of epididymal white adipose tissue (eWAT). Flow cytometry analysis of the SVF from eWAT showed that VLX-1005 treatment decreased the myeloid cell (CD11b+) population (**Figure 4E**). Using RT-PCR, we quantified mRNA expression of pro-inflammatory cytokines in the SVF and found significantly reduced expression of inflammatory markers, tumor necrosis factor α (TNFα) and arginase, with VLX-1005 treatment, and trends toward reduction of interleukin (IL)-1β, IL-6, or IL-10 (**Figure 4F**). These data suggest that 12-LOX inhibition with VLX-1005 reduces macrophage infiltration and associated inflammation in adipose tissue.

**Figure 4:**
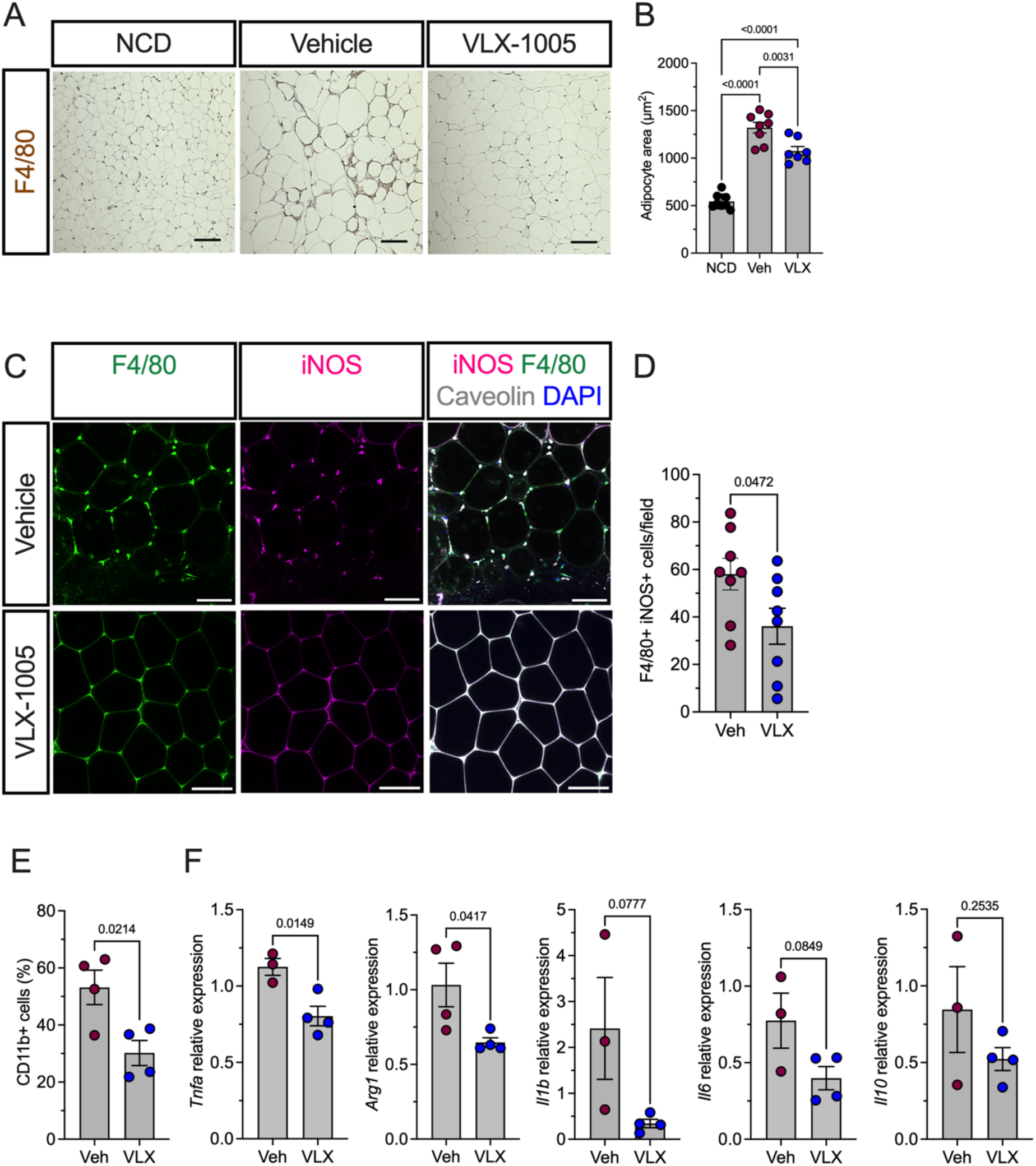
VLX-1005 treatment reduces infiltration of macrophages into pancreatic islets and adipose tissue. (A) Representative images from adipose tissue of *B6.hALOX12* mice on NCD treated with vehicle or HFD concurrently treated with vehicle or VLX-1005 showing immunohistochemical labeling for F4/80. Scale bars are 100 µm. (B) Adipocyte area of adipose tissue from *B6.hALOX12* mice on NCD treated with vehicle or HFD concurrently treated with vehicle or VLX-1005 (N=7-8 per group). (C) Representative images from whole-mounted adipose tissue of *B6.hALOX12* mice on HFD concurrently treated with vehicle or VLX-1005. Labeling for nuclei (blue), F4/80 (green), iNOS (magenta), and caveolin (gray). Scale bars are 100 µm. (D) Quantification of iNOS+ F4/80+ cells (N=8 per group). (E) Flow cytometry of the SVF from adipose tissue of *B6.hALOX12* mice on HFD concurrently treated with vehicle or VLX-1005 (N=4 per group). Cells were labeled for and sorted on CD11b. (F) Quantitative RT-PCR analysis of RNA from the SVF from adipose tissue. Data presented as mean with SEM. Single data points represent individual animals unless otherwise noted. Statistical significance determined by two-tailed Student’s t-testing.

### Lipoxygenase signaling in macrophages contributes to macrophage infiltration in both pancreatic islets and adipose tissue

Because inhibition of 12-LOX with VLX-1005 improves both islet and peripheral tissue parameters associated with macrophages during obesity, we asked if the effects of 12-LOX inhibition might be mediated through the action of the enzyme specifically in macrophages. In this regard, prior studies have established the impact of KO in whole-body, pancreas, and adipose tissue (10–13) but not macrophages. We crossed *B6.Alox15^fl/fl^* mice to *B6.Lyz2-Cre* mice to generate a myeloid-specific KO of *Alox15* (*B6.Alox15^Δmyel^*). Eight-week-old male *B6.Alox15^Δmyel^* mice had similar body weight, glucose tolerance, and adipocyte area as littermate control mice (**Supplemental Figure S2A–C**), implying that the constitutive loss of *Alox15* in myeloid cells did not affect metabolic parameters under normal chow conditions.

Male *B6.Alox15^Δmyel^* mice and their littermate controls were fed a HFD beginning at 8 weeks of age for 15 weeks in total (**Figure 5A**). Body weight, fat mass, and lean mass were identical between the control and *B6.Alox15^Δmyel^* mice (**Figure 5B–C**). Random-fed blood glucose values were also identical between the two groups (**Figure 5D**). A GTT after 8 weeks of HFD was indistinguishable between control and *B6.Alox15^Δmyel^* mice (**Figure 5E**). An ITT after 10 weeks of HFD also did not show differences between the two groups (**Figure 5F**). Analysis of pancreatic tissue from control and *B6.Alox15^Δmyel^* mice showed no difference in β cell mass (**Figure 5G**), however, there was reduced islet immunostaining for the β cell dedifferentiation marker ALDH1A3 in the *B6.Alox15^Δmyel^* mice (**Figure 5H**), similar to the VLX-1005-treated *B6.hALOX12* HFD-fed mice.

**Figure 5:**
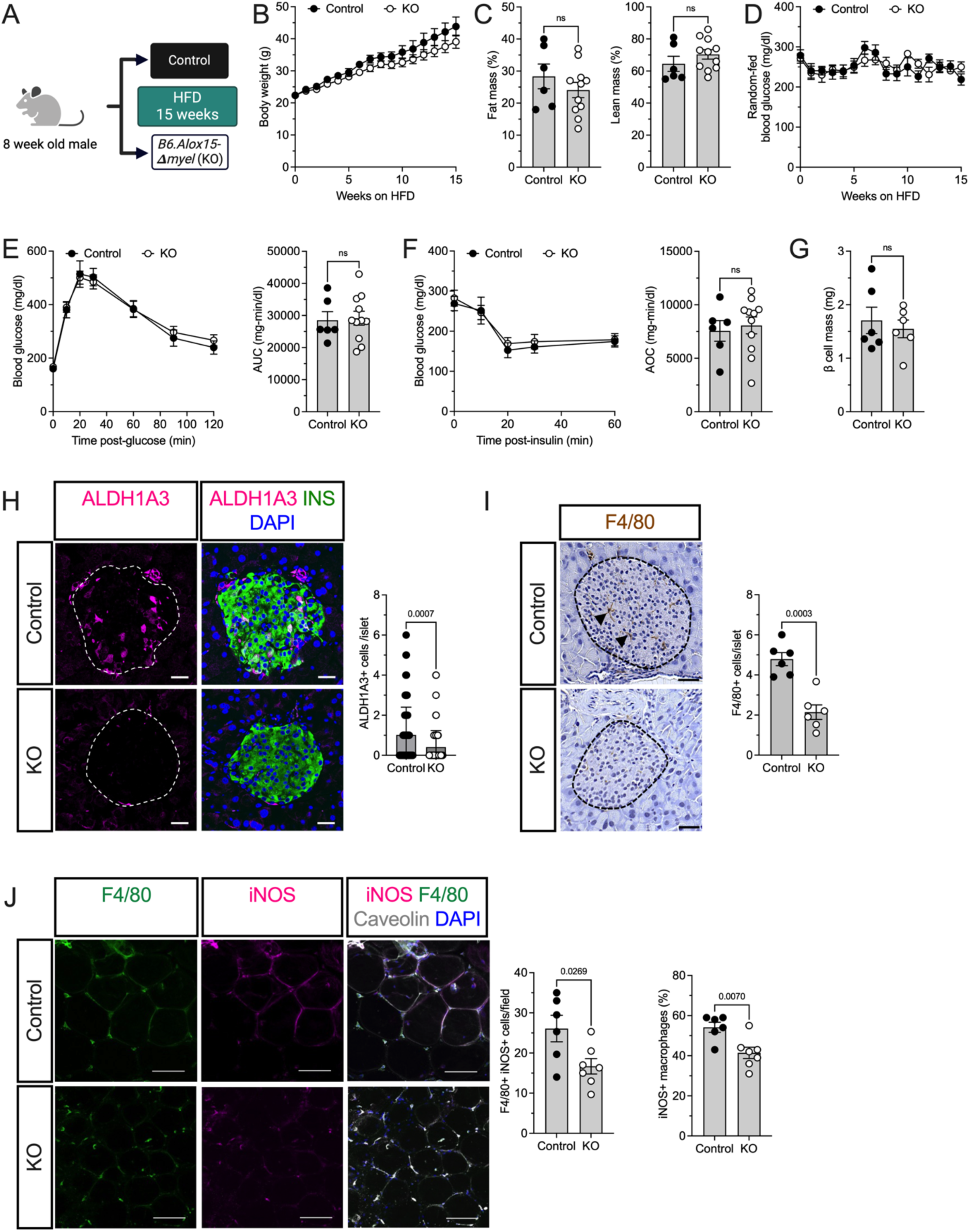
*Alox15* knockout in macrophages (*B6.Alox15^Δmyel^*or KO) does not change glucose homeostasis but reduces β cell dedifferentiation as well as macrophage infiltration in islets and adipose tissue. (A) Schematic depicting the experimental design. *B6.Alox15^Δmyel^* mice and littermate controls were treated with HFD for a total of 15 weeks (N=6 for control and N=11 for KO). (B) Body weight. (C) Fat and lean mass at endpoint. (D) Random-fed blood glucose measurements. (E) Blood glucose measurements from GTT at 8 weeks (*left panel*) and AUC (*right panel*). (F) Blood glucose measurements from ITT at 10 weeks (*left panel*) and AOC (*right panel*). (G) Quantification of β cell mass. (H) Representative images of pancreata with labeling for nuclei (blue), insulin (green), and ALDH1A3 (magenta) (*left panel*) and associated quantification (*right panel*) (N=80-88 islets per group). Scale bars are 20 µm. (I) Representative images of pancreata with immunohistochemical labeling for macrophages with F4/80 (arrowheads) (*left panel*) and associated quantification (*right panel*) (N=6 mice per group). Scale bars are 50 µm. (J) Representative images of whole-mounted adipose tissue labeled for nuclei (blue), F4/80 (green), iNOS (magenta), and caveolin (gray) (*left panel*). Quantification of total F4/80+ iNOS+ cells/field as well as iNOS+ cells as a percentage of F4/80+ cells (*right panel*) (N=6-7 per group). Scale bars are 100 µm. Data presented as mean with SEM. Single data points represent individual mice unless otherwise noted. Statistical significance determined by two-tailed Student’s t-testing.

We next assessed macrophage accumulation in pancreatic tissue using immunohistochemistry for F4/80, which showed decreased F4/80+ cells per islet (**Figure 5I**). In adipose tissue, we observed no change in adipocyte area in the *B6.Alox15^Δmyel^* mice (**Supplemental Figure 2C**). However, whole-mount immunofluorescence of F4/80+ and iNOS+ cells in adipose tissue showed a reduction in mice of total F4/80+ iNOS+ cells as well as iNOS+ cells as a percentage of all F4/80+ cells in *B6.Alox15^Δmyel^* mice (**Figure 5J**). Collectively, these data suggest the possibility that 12-LOX expression in macrophages drives infiltration of pro-inflammatory macrophages into both pancreatic and adipose tissue under HFD conditions, but without significant impact on glucose homeostasis.

## DISCUSSION

In this study, we used a translationally relevant human gene replacement mouse model of obesity and T2D to examine the effects of the 12-LOX inhibitor VLX-1005 on β cells and adipose tissue. Additionally, we established complementary models, including zebrafish and myeloid-specific *Alox15* KO mice, to investigate the impact of VLX-1005 treatment and the cell-type-specific role of 12-LOX in obesity-related inflammation. We hypothesized that inhibition of human 12-LOX with VLX-1005 would recapitulate the previous findings of whole-body KO of 12-LOX in mice, improving glucose homeostasis by decreasing both β cell dysfunction and adipose tissue inflammation. Collectively, our data show that VLX-1005 treatment improves glucose homeostasis, reduces β cell dedifferentiation, and alleviates macrophage-mediated inflammation in both pancreatic islets and adipose tissue.

Prior studies in rodent models and tissues from human T2D donors have shown an increase in islet-associated macrophages, and mechanistic studies further demonstrate increased release of cytokines from islets (18,19,27,28). In addition to known roles in chemotaxis, local action of cytokines such as TNFα and IL-1β have been shown to negatively impact β cell function directly (18,29,30). There is also evidence of reciprocal proinflammatory signaling between β cells and macrophages through palmitate and toll-like receptor 4 (TLR4) which leads to infiltration of islets by macrophages and β cell dysfunction (18). We show that 12-LOX inhibition with VLX-1005 treatment reduces macrophage numbers in the islets of both zebrafish and mice. In the context of these known mechanisms, this finding suggests VLX-1005 treatment may contribute to decreased proinflammatory signaling in islets and improved β cell function from decreased β cell exposure to macrophages.

We also observed decreased β cell dedifferentiation with VLX-1005 treatment by immunolabeling for ALDH1A3. In mice, ALDH1A3 expression is associated with β cell failure and reduced glucose-stimulated insulin secretion, mitochondrial dysfunction, and progenitor-like features (17). Analysis of pancreatic tissue from donors with T2D shows that β cell dedifferentiation is increased compared to control donors without T2D, and that this finding is correlated with decreased glucose-stimulated insulin secretion and endocrine cell maturity (16,31,32). The decrease in ALDH1A3 with VLX-1005 treatment that we observe in our study, therefore, suggests improved β cell health and potential capacity to meet increased insulin secretory needs.

Obesity-associated inflammation in adipose tissue contributes to insulin resistance through multiple mechanisms mediated by both adipocytes and macrophages (23,33,34). These mechanisms include changes in adipocyte production of free fatty acids, lipopolysaccharide, adipokines, and cytokines, including TNFα, IL-1β, IL-6, IL-8, and C-X-C motif ligand 1 (CXCL1) (35–39), which lead to proinflammatory macrophage recruitment and polarization of resident adipose tissue macrophages. These proinflammatory macrophages contribute to inflammatory crosstalk with adipocytes, further macrophage recruitment, and ultimately downstream changes in insulin receptor signaling and insulin resistance (24,33,40,41). Prior studies in rodent models have identified C-C motif ligand 2 (CCL2; also known as monocyte chemotactic protein 1 or MCP-1) as having a key role in both polarization and recruitment of macrophages through action on the receptors C-C motif chemokine receptor 2 (CCR2) and CCR5 (42–44). Macrophages treated with 12-HETE or overexpressing *Alox15* show increased expression of mRNA for *Ccl2* as well as *Il6* and *Tnfa* (45), and analysis of adipose tissue in *Alox15* KO mice showed decreased CCL2 expression and reduced macrophage infiltration (10). In our studies, we show that 12-LOX inhibition with VLX-1005 treatment reduced macrophage infiltration and cytokine expression in adipose tissue, and this may relate in part to reduced secretion of CCL2. We also show an increase in insulin sensitivity with VLX-1005 treatment, similar to whole-body 12-LOX KO (11) but in contrast with the pancreas-specific 12-LOX KO, which did not result in an improvement in insulin sensitivity (12). This findings uggests that 12-LOX activity in macrophages contributes to insulin resistance.

We investigated a contribution of 12-LOX in macrophages by studying mice with deficient 12-LOX in macrophages. The *B6.Alox15^Δmyel^* mice revealed no changes in glucose homeostasis but did show decreased β cell dedifferentiation and reduced macrophage infiltration in both islets and adipose tissue. These findings agree with the previously identified role for the products of 12-LOX enzyme activity, namely 12-HETE, in mediating leukocyte migration (46,47). 12-HETE may act by binding to receptors known to stimulate chemotactic responses. 12-HETE has a low affinity for the leukotriene B4 receptor 2 (BLT2) (48) and a high affinity for G-protein coupled receptor 31 (GPR31) (49). Our group has previously investigated the role of BLT2 in macrophage migration in *BLT2* KO mice, showing decreased migration of macrophages to the peritoneum and liver in response to lipopolysaccharide (LPS) treatment without associated changes in macrophage polarization (50). While we have limited data on the role of GPR31 in mammals in this context, prior work in zebrafish has shown that the orthologue *gpr31* is expressed in pancreas and its knockdown phenocopies loss of 12-LOX activity while also impairing pancreatic organogenesis (51). Finally, it is possible that loss of 12-LOX activity directly impacts proinflammatory signaling in macrophages in addition to chemotaxis. Consistent with this possibility, macrophages isolated from NOD mice in which human *ALOX12* replaced mouse *Alox15* show reduced interferon gene responses after inhibition of 12-LOX with VLX-1005 treatment (14). Altogether, our findings suggest that the enzymatic activity of 12-LOX impacts macrophage migration into adipose tissue and islets, and that improvements in glucose homeostasis with 12-LOX inhibition are a consequence of: (1) decreased macrophage infiltration into islets; (2) decreased proinflammatory action of 12-HETE produced in pancreatic tissue by macrophages or β cells; or (3) increased peripheral insulin sensitivity through reduced macrophage infiltration in adipose tissue.

There are some limitations to our study. First, there are known differences in tissue-specific expression of *Alox15* in mice compared to *ALOX12* in human, as well as sequence differences between orthologous LOX family members. Our mouse model replaces mouse *Alox15* with human *ALOX12* globally, and it is known that mouse *Alox15* is also expressed in other leukocytes, epithelial cells, reticulocytes, and smooth muscle (9). Therefore, we cannot rule out that the effects of VLX-1005 inhibition are related to *ALOX12* enzyme expression in cells other than β cells or macrophages. Second, the two study designs we utilized may not fully reflect the chronicity of obesity and T2D. It is possible that the negative impact of proinflammatory signaling through the products of 12-LOX may accumulate over time, and that longer treatment with VLX-1005 could have a greater effect size on insulin sensitivity and glucose homeostasis parameters. Third, we utilized a dose of VLX-1005 (30 mg/kg/d) that was optimized in a type 1 diabetes prevention study (14), and it may be that the optimal dose regimen in the setting of obesity and T2D requires further titration.

In summary, our study establishes a platform to study the inhibition of 12-LOX in the setting of obesity and dysglycemia. Inhibition of 12-LOX in our human gene replacement mouse model recapitulates some but not all features of prior 12-LOX KO studies, highlighting differences between human LOX enzymes and those of other species as well as the need to assess pharmacologic inhibitors using human-relevant model systems. Our mouse model also provides a platform to investigate the role of 12-LOX in other metabolic–inflammatory diseases, such as atherosclerosis, endothelial dysfunction, and metabolic dysfunction-associated steatotic liver disease. Lastly, our study underscores the potential of the 12-LOX inhibitor VLX-1005 as a therapeutic agent in managing obesity and dysglycemia by targeting both β cell factors and peripheral inflammation associated with visceral adiposity.

## MATERIALS AND METHODS

### Animals

All experiments involving zebrafish and mice were performed at the University of Chicago and the procedures were conducted according to protocols approved by the University of Chicago Institutional Animal Care and Use Committee.

The generation of the *B6.hALOX12* mouse line has been previously published (14). To generate *Lyz2-Cre;Alox15^fl/fl^* mice on the *C57BL/6J* background (*B6.Alox15^Δmyel^*), *B6.Lyz2-Cre* (Jackson Laboratory; strain #004781) were crossed *to B6.Alox15^fl/fl^* mice (Jackson Laboratory; strain #031835). All mice were kept under pathogen-free housing conditions with standard light:dark (12:12) cycles and fed *ad lib* NCD until initiation of the HFD treatments as described below.

Transgenic zebrafish (*D*. *rerio*) expressing GFP under the macrophage-specific promoter *mpeg*, *Tg(mpeg:eGFP)gI22* were obtained from the Zebrafish International Resources Center (catalog #ZL9940). Zebrafish were kept at 28.5 °C under standard laboratory conditions at our institution.

### Glucose homeostasis testing

Intraperitoneal GTT were performed in mice after overnight fasting (16 h). Mice were intraperitoneally injected with glucose at a dose of 1 g/kg lean body mass and blood glucose levels were measured at 0, 10, 20, 30, 60, 90, and 120 min after glucose injection by glucometer (AlphaTrak).

For glucose-stimulated insulin secretion testing, mice were injected with 2 g/kg lean body mass and blood was collected at 0 and 10 min. Insulin was analyzed using ultrahigh sensitivity insulin enzyme-linked immunosorbent assay (Crystal Chem; 90080). HOMA-IR values were calculated using fasting glucose (mg/dl) prior to glucose injection multiplied by fasting insulin (µIU/ml) divided by 405.

Intraperitoneal ITT were performed in mice after fasting for 2 h. Mice were intraperitoneally injected with insulin at a dose of 0.75 U/kg or 1.5 U/kg body weight and blood glucose was measured at 0, 15, 30, 45, and 60 min after insulin injection.

### Treatment with VLX-1005

VLX-1005 spray-dried dispersion (SDD) was obtained from Veralox Therapeutics, Inc.

Preparation and characterization of VLX-1005 SDD was as previously described (14). VLX-1005 or vehicle was given to all mice via oral gavage.

### HFD pretreatment before VLX-1005 experiments

Male *B6.hALOX12* mice at 8 weeks of age (n=23) were started on HFD (60% of total calories from fat; Research Diets; D12492) and maintained on this diet for 8 weeks, prior to dividing into treatment groups: vehicle (n=11) and 30 mg/kg/day of VLX-1005 (n=12). A GTT was performed at 12 weeks. An ITT was performed at 14 weeks after which animals were sacrificed, and tissues were collected for analysis.

### HFD and VLX-1005 concurrent treatment experiments

Male *B6.hALOX12* mice at 8 weeks of age were started on NCD (6% total calories from fat, Inotiv; #2918) or HFD (60% of total calories from fat; Research Diets; D12492) and split into vehicle (n=19) or 30 mg/kg/day of VLX-1005 (n=18). GTT was performed after 8 weeks of diet. ITT was performed after 10 weeks of diet, and the animals were subsequently sacrificed and tissues collected for analysis. A subset of these animals in both vehicle (n=4) and treatment (n=4) groups were injected with 3 U/kg insulin 5 minutes before sacrifice for Western blot analysis of AKT and phosphorylated AKT as described below.

### B6.Alox15^Δmyel^ experiments

Male *B6.Alox15^Δmyel^* mice at 8 weeks of age (n=11) and littermate controls (N=6) were started on HFD (60% of total calories from fat; Research Diets; D12492). A GTT was performed at the start of the study and 8 weeks after diet initiation. An ITT was performed at 10 weeks after diet initiation. The animals were sacrificed at 15 weeks for further tissue analysis.

### Zebrafish experiments

Embryos were collected at spawning and incubated at 28.5 °C in standard maintenance media (0.1% instant ocean salt, 0.0075% calcium sulfate). After gastrulation, 0.003% 1-phenyl-2-thiourea (Acros; #207250250) supplementation in maintenance media was used to prevent pigmentation in all embryos and larvae. At 4 and 6 dpf, zebrafish were treated with a solution of 5% whole chicken egg for 3 h and returned to maintenance media after the end of each treatment. Starting on 4 dpf, vehicle (dimethyl sulfoxide) or VLX-1005 (10 µM) was added to either maintenance media or chicken egg solutions as needed for continuous treatment until 7 dpf, at which point larvae were fixed in 2.5% formaldehyde in buffered saline at 4 °C overnight and immunolabeled as previously described (52). We labeled whole-mount zebrafish with primary anti-insulin (Abcam; #ab210560), secondary anti-rabbit (Invitrogen; #A10042), and 4’,6-diamidino-2-phenylindole (DAPI; Thermo Fisher) for nuclei. After immunolabeling, larvae were mounted on slides in antifade mounting medium (Vector Labs). An A1 confocal microscope (Nikon) was used to capture immunofluorescence images. The number of GFP+ cells within islets was assessed manually in Fiji (53).

### Immunolabeling and β cell mass

Tissues from animals were fixed using 4% paraformaldehyde for 4 hours for pancreas, 24-48 hours for liver, and 72 hours for adipose tissue. These tissues were subsequently embedded in paraffin and sectioned with a thickness of 5 µm. For analysis of pancreas, we used three sections per mouse with each section spaced 100 µm apart.

For immunohistochemistry, tissue sections were initially immunolabeled with primary anti-insulin (ProteinTech; 15848-1-AP; 1:200), anti-glucagon (Abcam; ab92517; 1:200), and anti-F4/80 (Sigma; D2S9R; 1:150). Secondary antibody immunolabeling was with conjugated anti-rabbit Ig antibody (Vector Laboratories). 3,3’-Diaminobenzedine (DAB) peroxidase substrate kit (Vector Laboratories) was used for detection per the manufacturer’s instructions. After immunolabeling, the tissue sections were counterstained with hematoxylin (Sigma). Images were acquired using a BZ-X810 microscope (Keyence). β cell mass was quantified by insulin-positive area and whole pancreas area.

For immunofluorescence analysis, tissue sections were stained with the following antibodies: primary anti-insulin (ProteinTech; 15848-1-AP; 1:200), anti-glucagon (Abcam; ab92517; 1:200), and anti-ALDH1A3 (Novus; NBP2-15339; 1:500). Highly cross-adsorbed AlexaFluor secondary antibodies (Thermo Fisher) were used at a dilution of 1:250. Tissue sections were stained with DAPI to label cell nuclei.

For whole-mount immunofluorescence of adipose tissue, adipose tissue was excised fresh from mice and placed in 1% paraformaldehyde for 1 hour. These tissues were subsequently stained with primary antibodies against iNOS (Novus, NBP1-33780), caveolin (Novus, NB100-615), and F4/80 (Abcam, ab6640) at a dilution of 1:100 and incubated at 4° C overnight. Tissues were subsequently immunolabeled with AlexaFluor secondary antibodies (Thermo Fisher) at a dilution of 1:250, stained with DAPI as above, and mounted on gasketed coverslips for imaging.

An A1 confocal microscope (Nikon) was used to capture immunofluorescence images. The number of cells positive for each marker were assessed manually unless otherwise noted. For analysis of whole-mounted adipose tissue images in *B6.hALOX12* mice, we also used the density counting workflow in the ilastik software package to train a classifier to identify and count F4/80+ and iNOS+ cells (54). For automated calculation of adipocyte geometry, we used Adiposoft software (55).

### Western blot analysis

Tissue was collected from animals in radio-immunoprecipitation assay (RIPA) buffer containing phosphatase and protease cocktails per the manufacturer’s instructions (Thermo Fisher). We used 40 µg of sample per lane and immunolabeled with primary anti-AKT (Cell Signaling; 9272; 1:1,000), phospho-AKT (Cell Signaling; 4060S; 1:1,000), and β-actin (Cell Signaling 3700; 1:1,000) and secondary donkey antibodies against rabbit and mouse (LICOR; 926-68073 and 926-32212; 1:10,000). Immunolabeled membranes were imaged on an Odyssey near infrared imaging system (LI-COR) and quantified using Fiji (53).

### Stromal vascular fractionation

Epididymal white adipose tissue from *B6.hALOX12* mice on HFD concurrently treated with vehicle and VLX-1005 was harvested and immediately placed in ice-cold phosphate buffered saline (PBS) containing 3% fetal bovine serum. The adipose tissue was minced into 2-4 mm pieces using scissors and transferred into C tubes (Miltenyi Biotec). The C tubes were then attached upside down onto the sleeve of the gentleMACS dissociator (Miltenyi Biotec) and run twice. The dissociated tissue was then filtered through a 100 μm strainer and centrifuged at 500 g for 5 minutes at 4°C. After centrifugation, the supernatant was removed using a vacuum filtration system to eliminate residual fats and lipids. The pellet was then resuspended in PBS.

### Flow cytometry

The stromal vascular fraction (SVF) was resuspended in stain buffer (BD Pharmingen, 554657) and incubated with a blocking solution containing anti-mouse CD16/CD32 (eBioscience; 14-0161-86) for 20 minutes on ice to inhibit Fc receptor binding. After blocking, the cells were washed with staining buffer and subsequently incubated for 20 minutes on ice with fluorochrome-conjugated antibodies for CD11b (Biolegend; 101225) and F4/80 (Biolegend; 123108) at a 1:1,000 dilution. Following antibody staining, cells were washed and analyzed using an Attune NxT Flow Cytometer (Thermo Fisher). Data analysis was conducted using FlowJo software (BD Biosciences). Antibody specificity was confirmed through control experiments, which included the omission of primary or secondary antibodies and gating to exclude background fluorescence.

### Quantitative RT-PCR analysis

RNA was isolated from the SVF utilizing the RNeasy Mini Kit (Qiagen) and cDNA was synthesized using the High-Capacity cDNA Reverse Transcription Kit (Applied Biosystems) following the manufacturer’s instructions. Quantitative real-time PCR (RT-PCR) was performed using the SensiFAST SYBR Lo-ROX Kit (Thomas Scientific) on a CFX Opus system (Bio-Rad) using the primers listed in **Supplemental Table S1**. Relative gene expression was determined by normalizing the average comparative threshold cycle (CT) values of each gene to *Hprt1* expression. The normalized expression levels were then calculated relative to the averaged vehicle controls using the ΔΔCT method.

### Sex as a biological variable

We used male mice alone as comparative data in the literature exists primarily for male mice. There is also known differential weight gain based on sex while on HFD with female mice gaining less weight than male mice (56).

### Statistical methods

All data shown are mean with error bars representing standard error of the mean (SEM). When comparing two conditions, two-tailed Student’s t-tests were performed. When comparing experiments with more than two conditions, one-way ANOVA was performed followed by Dunnett’s or Tukey’s correction for multiple comparisons to determine pairwise differences between groups. Differences between groups were considered statistically significant for P values <0.05. Data analyses were performed using GraphPad Prism 10 software.

## AUTHOR CONTRIBUTIONS

JLN, MBB, DJM, RNA, SAT, and RGM conceptualized the research. KBK, TN, KF, JEW, SP, AC, IC, and SAT performed experiments and analyzed the data. SAT and RGM provided project supervision. KBK, TN, SAT, and RGM wrote the original manuscript draft. All authors contributed to discussion, edited, and approved the final version of the manuscript.

## ACKNOWLEDGEMENTS

This work was supported in part by National Institutes of Health grants R03 TR003381 (to SAT and RGM), R41 DK122917 (to RGM and DJM), R01 DK10558 (to RGM), T32 DK007011 (to KBK), and an investigator-initiated award from Veralox Therapeutics (to SAT and RGM). This study utilized Diabetes Center core resources supported by National Institutes of Health grant P30 DK020595 (to the University of Chicago) and utilized services of the University of Chicago Human Tissue Resource Center. Schematics were created with BioRender.

## SUPPLEMENTAL FIGURES

**Supplemental Figure S1:**
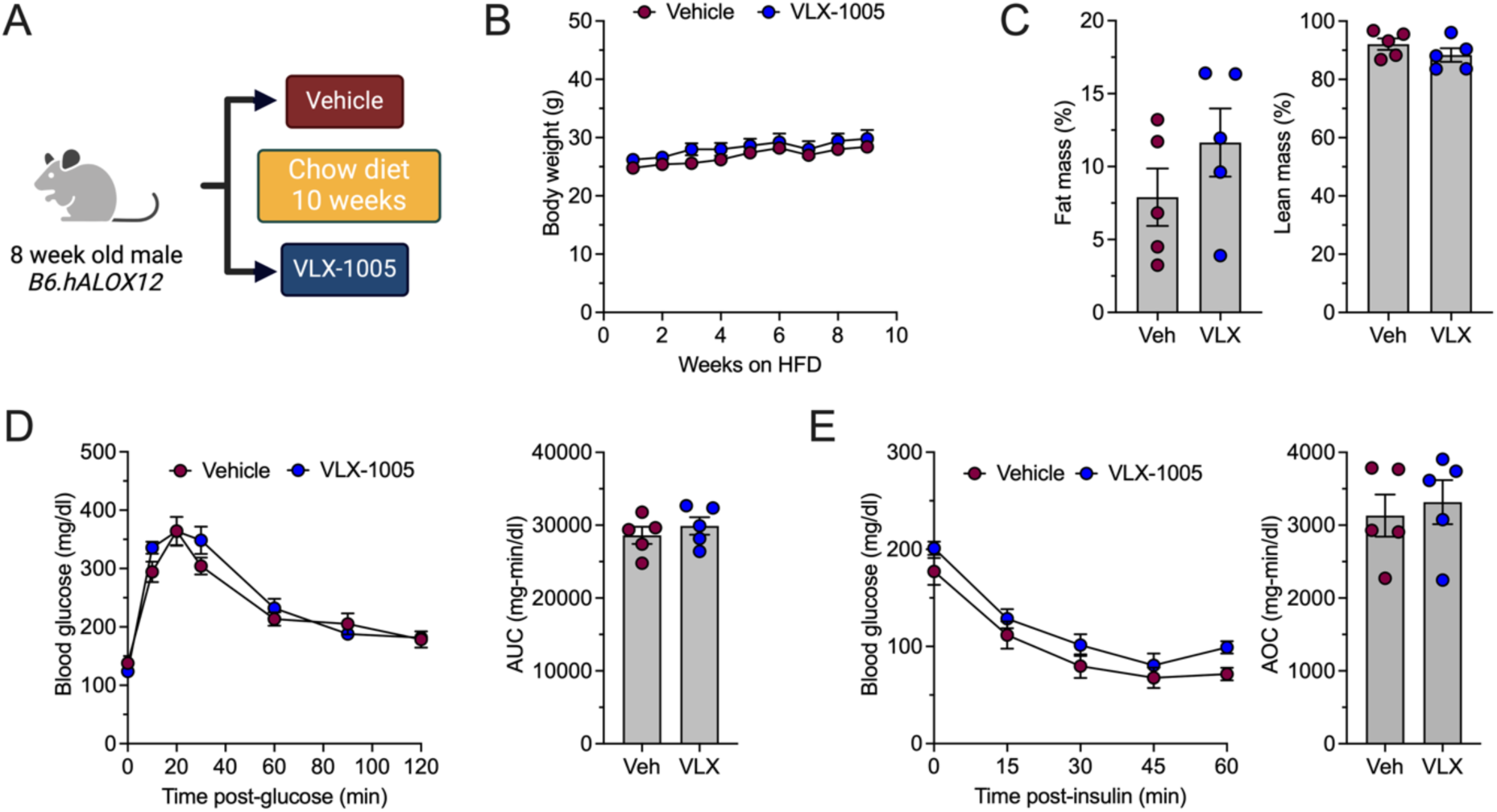
*B6.hALOX12* mice on NCD treated with VLX-1005 show no changes in the size of the adipose compartment or glucose homeostasis. (A) Schematic depicting the experimental design. *B6.hALOX12* mice were treated with NCD concurrently with either vehicle or VLX-1005 for 10 weeks (N=5 per group). (B) Body weight. (C) Fat and lean mass. (D) Blood glucose measurements from GTT at 8 weeks (*left panel*) with AUC values (*right panel*). (E) Blood glucose measurements from ITT at 10 weeks (*left panel*) with AOC values (*right panel*). Data presented as mean with SEM. Single data points represent individual mice. Statistical significance determined by two-tailed Student’s t-testing.

**Supplemental Figure S2:**
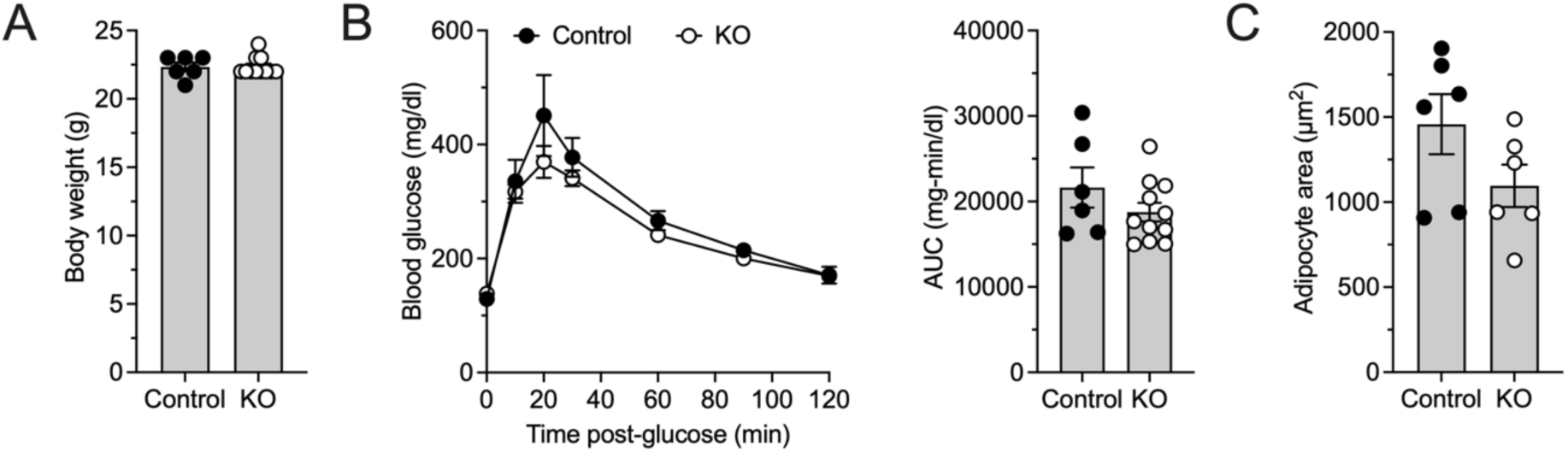
Additional data for *B6.Alox15^Δmyel^* (KO) and littermate control mice. (A) Body weight prior to initiation of HFD. (B) Blood glucose measurements from GTT at start of study prior to initiation of HFD (*left panel*) and AUC (*right panel*) (N=6 for controls and N=11 for KO). (C) Adipocyte area (N=6-7 per group). Data presented as mean with SEM. Single data points represent individual mice. Statistical significance determined by two-tailed Student’s t-testing.

## SUPPLEMENTAL TABLES

**Supplemental Table S1:**
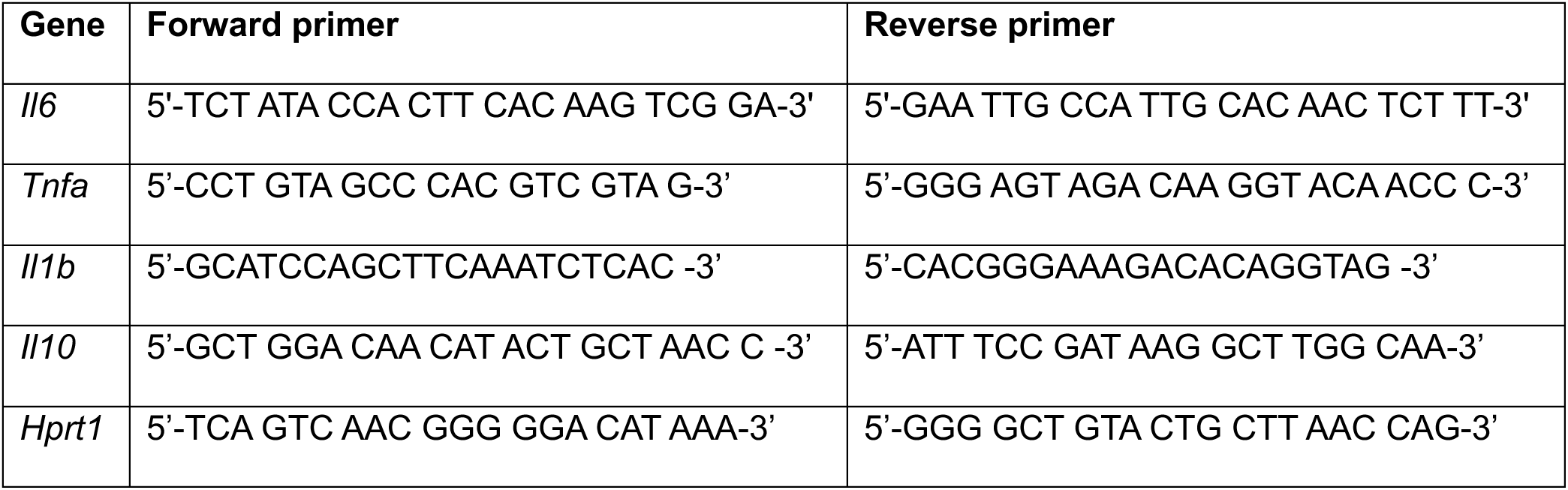
Primer pairs for quantitative RT-PCR.

